# Arrhythmogenesis as the failure of repolarization

**DOI:** 10.1101/2020.07.02.185694

**Authors:** Stewart Heitmann, Jamie I Vandenberg, Adam P Hill

## Abstract

Contemporary theories of cardiac fibrillation typically rely on the emergence of rotors to explain the transition from regular sinus rhythm to disordered electrophysiological activity. How those rotors spontaneously arise in the absence of re-entrant anatomical circuits is not fully understood. Here we propose a novel mechanism where arrhythmias are initiated by cardiac cells that fail to repolarize following a normal heartbeat. Those cells subsequently act as a focal ectopic source that drive the ensuing fibrillation. We used a simple computational model to investigate the impact of such cells in both homogeneous and heterogeneous excitable media. We found that heterogeneous media can tolerate a surprisingly large number of abnormal cells and still support normal rhythmic activity. At a critical limit the medium becomes chronically arrhythmogenic. Numerical analysis revealed that the critical threshold for arrhythmogenesis depends on both the strength of the coupling between cells and the extent to which the abnormal cells resist repolarization. Arrhythmogenesis was also found to emerge first at tissue boundaries where cells naturally have fewer neighbors to influence their behavior. These findings may explain why atrial fibrillation typically originates from the cuff of the pulmonary vein.

**Author summary:** Cardiac fibrillation is a medical condition where normal heart function is compromised as electrical activity becomes disordered. How fibrillation arises spontaneously is not fully understood. It is generally thought to be triggered by premature depolarization of the cardiac action potential in one or more cells. Those premature beats, known as early-afterdepolarizations, subsequently initiate a self-sustaining rotor in the otherwise normal heart tissue. In this study, we propose an alternative mechanism whereby arrhythmias are initiated by cardiac cells that fail to repolarize of their own accord but still operate normally when embedded in functional heart tissue. We find that such cells can act as focal ectopic sources under appropriate conditions of inter-cellular coupling. Moreover, cells on natural tissue boundaries are more susceptible to arrhythmia because they are coupled to fewer cells. This may explain why the pulmonary vein is often implicated as a source of atrial fibrillation.

## Introduction

Cardiac arrhythmias, which impair the heart’s ability to pump blood effectively, are a major cause of morbidity [1] and mortality [2]. Arrhythmic activity is underpinned by re-entrant rotors of electrical activity in the myocardium [3–6]. However, the processes by which rotors are initiated and subsequently breakup into disordered activity are not well understood. This lack of knowledge is reflected in the paucity of effective treatment options for patients with cardiac arrhythmias.

Contemporary understanding of arrhythmogenesis revolves around the concepts of triggers and a vulnerable substrate. The trigger for an arrhythmia is often attributed to spontaneous ectopic activity, such as early after-depolarizations (EADs). For example, the majority of cases of atrial fibrillation are thought to be triggered by ectopic activity originating at the cuff of the pulmonary vein [7,8]. However, theoretical models suggest that EADs occurring in individual myocytes will fail to initiate sustained propagating activity because of the damping effect of electrotonically coupled tissue [9]. Rather, ectopic beats will only initiate arrhythmias if they land in a vulnerable substrate. At least three factors can contribute to making the myocardium more vulnerable to sustaining an arrhythmia. First, topological defects in the heart that form anatomical re-entry circuits, such as AV nodal accessory pathways [10]. Second, reduced intercellular connectivity, as for example occurs in stretched atria with diffuse interstitial fibrosis [1]. And third, heterogeneity in the refractoriness of cardiac cells, which is also thought to be key to the breakdown of orderly waves into multiple wavelets [5,9,11,12].

The duration of the refractory period in heart cells is dependent on the duration of the action potential. In the normal heart, there are regional variations in action potential duration, for example the epicardial to endocardial gradients in the ventricle [13] and systematic regional differences in the atria [14]. It is generally thought that the repolarization duration of individual cells, whilst changing gradually between regions, all fall on a continuum within a relatively narrow physiological range [15,16]. However, as repolarization of the cardiac action potential represents a delicate balance between inward and outward currents, small perturbations to individual currents can have significant effects on repolarization. In a recent study of individual atrial myocytes, from both rabbit and human, Kettlewell and colleagues [17] showed that widening the voltage window for calcium channel reactivation by as little as 6 mV, can result in extreme changes in the action potential duration and in some instances the cells fail to repolarize. Such cells are said to be ‘bistable’ because they are capable of residing in either the normal resting state or at an elevated voltage state [17].

We hypothesized that if there are heart cells that could exhibit bistability that they could provide both a substrate and a trigger for arrhythmogenesis. The extreme variation in repolarization heterogeneity could contribute to the breakdown of orderly wave propagation that occurs in fibrillation. Furthermore, they could also be good candidates for initiating fibrillation because they could remain in the depolarized state well beyond the refractory period of their neighbors. In this study we undertook a theoretical assessment of how bistable cells might contribute to the genesis of fibrillation by modeling heart tissue as an excitable medium of FitzHugh-Nagumo cells [18,19] that were configured to have either mono- or bistable membrane potentials. We then investigated how the behavior of bistable cells was influenced by electrical coupling, structural features of cardiac tissue, dimensionality of connectivity and heterogeneity of repolarization duration.

## Results

In order to examine the impact of bistable cells in an electrically coupled myocardium, we modeled the heart as a two-dimensional sheet of excitable cells with generalized FitzHugh-Nagumo [18,19] dynamics,

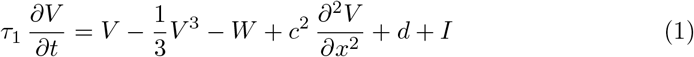

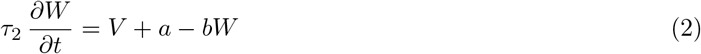

where *V*(*x,t*) represents the membrane potential of the cell at position *x*∈*R*^2^. The recovery variable *W*(*x, t*) is an abstract representation of the repolarizing currents that return the membrane potential to rest. Parameter *I*(*x,t*) is a spatiotemporal stimulus that is applied to the medium to initiate a propagating wave. Parameters *a, b* and *d* are constants that dictate the dynamics of the individual cells. Parameter *c*^2^ represents the strength of the electrical coupling between adjacent cells. Equations (1–2) encapsulate the fundamental character of the heart as an excitable medium without the computational burden of excessive anatomical or molecular detail.

Excitability is best understood by analyzing the phase plane for an isolated cell (Figures 1A,D; Equations 7–8 in Methods). The phase plane describes how the states of *V* and *W* evolve with respect to each other. The nullclines (green) indicate those points in the phase plane where the time derivatives of each state variable are zero. Equilibrium points exist where the nullclines intersect. In this case, the nullcline for 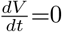 is the cubic,

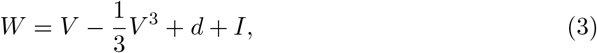

and the nullcline for 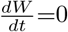 is the straight line,

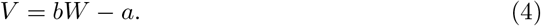

**Fig 1.**
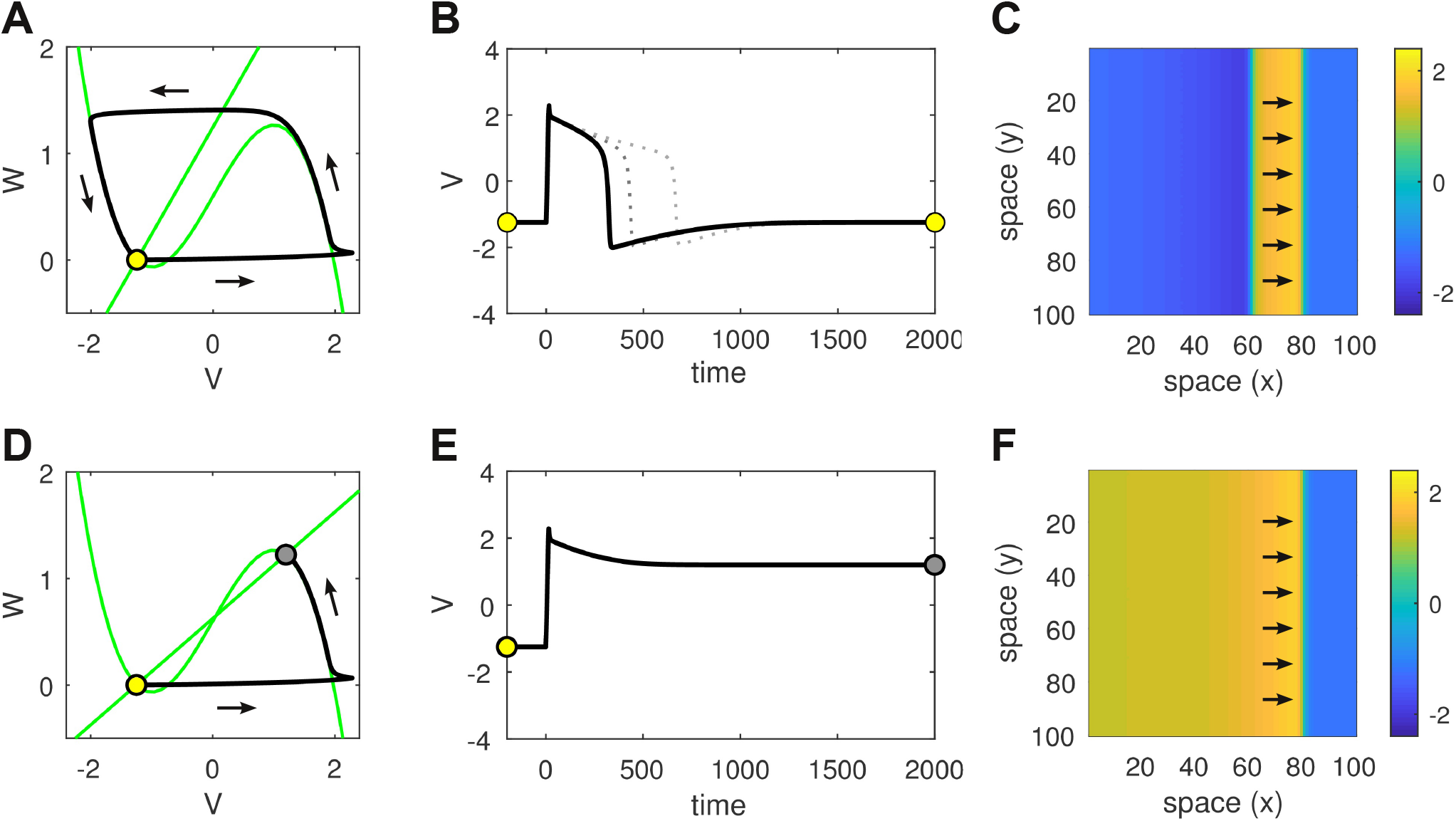
Comparison of monostable versus bistable dynamics in both the isolated cell and the homogeneous spatial medium. **(A)** Phase plane of the isolated cell in the monostable regime (*b*=1). The nullclines are shown in green. The single fixed point (open circle) is stable. It represents the resting membrane potential. The system can be perturbed from rest by a brief injection current (*I* = 2 for 15 ms) that initiates a trajectory (black line) which makes a large excursion in phase space before returning to rest. That excursion corresponds to the action potential. **(B)** Time plot of the action potential in the monostable regime. Solid line for *b* = 1. Dotted lines for *b* = 1.5 and *b* = 1.75. **(C)** Snapshot of the corresponding traveling wave in a homogeneous sheet of 100 × 100 monostable cells with coupling parameter *c*=1. Arrows indicate the direction of travel. Color indicates the membrane potential. **(D)** Phase plane of the isolated cell in the bistable regime (*b*=2). Perturbing the resting state (open circle) induces a transition to the up-state (filled circle). **(E)** Time plot of that transition. **(F)** The corresponding traveling front in the spatial medium.

We configured the parameters *a* and *d* so that the nullclines intersected near the left-hand knee of the cubic nullcline. Such an arrangement is a known requirement for excitability in the FitzHugh-Nagumo model and excitable systems in general [20]. More specifically, we chose *a*=1.25 and *d*=0.6 so that the equilibrium point (*V*=−1.25, *W*=0) was independent of parameter *b* because *W*=0.

Parameter *b* dictates the slope of the *W* nullcline (equation 4) which hereby pivots on the equilibrium point for our choice of *a* and *d*. For the case of *b*=1, the nullclines intersect exactly once and the system is *monostable* (Figure 1A). In this regime, an action potential can be elicited in the resting cell by injecting it with a brief injection current (Figure 1B). The time and space constants (*τ*_1_=10, *τ*_2_=400, *dx*=1) were chosen so the shape of the action potential best resembled those of cardiac cells. This action potential propagates as a planar wave in a homogeneous medium (Figure 1C). The propagation speed is predominantly proportional to the coupling parameter c. It is also effected to a lesser extent by the threshold of excitability defined by parameter *d*.

Small increases in *b* prolong the action potential (dotted lines in Figure 1B) by increasing the rate at which the recovery variable decays. In the context of cardiac cells, it is akin to shortening the lifetime of the repolarizing currents. However there is a limit to how much the recovery variable can be shortened before repolarization dramatically fails. Figure 1D shows the case of *b*=2 where the nullclines intersect at three points in the phase plane. Two of these fixed points are stable (marked by circles) and the other is unstable. The lower stable fixed point (open circle) corresponds to the same resting membrane potential (*V*=−1.25) as before. Whereas the upper stable fixed point (filled circle) represents a newly created high-voltage state (*V*=1.2) which we refer to as the *up-state*.

Crucially, the resting state and the up-state co-exist for the same choice of parameters, hence the cell is *bistable*. It can be transitioned from the resting state into the up-state using the same brief injection current as before (Figure 1E). In the spatial medium, that transition from resting state to up-state propagates as a traveling front which ultimately recruits the entire medium (Figure 1F).

### Stability analysis

To better understand the onset of bistability in the isolated cell, we used numerical continuation to follow the steady-state membrane potential over a range of values of *b* (Figure 2A). The resting state (*V* = − 1.25) was found to be stable (solid line) for all values of *b* that we investigated. Furthermore, it is the only solution that exists for *b*<1.64 meaning that regime is monostable. At *b*=1.64 an additional pair of unstable fixed points (dashed lines) emerge via a saddle-node (SN) bifurcation. Geometrically, this occurs when the linear nullcline comes into contact with the upper branch of the cubic nullcline. However the newly emerged fixed points are both unstable so the dynamical regime is still monostable at that point. Bistability does not emerge until *b*=1.76 where the up-state becomes stable via a supercritical Hopf bifurcation (HB).

**Fig 2.**
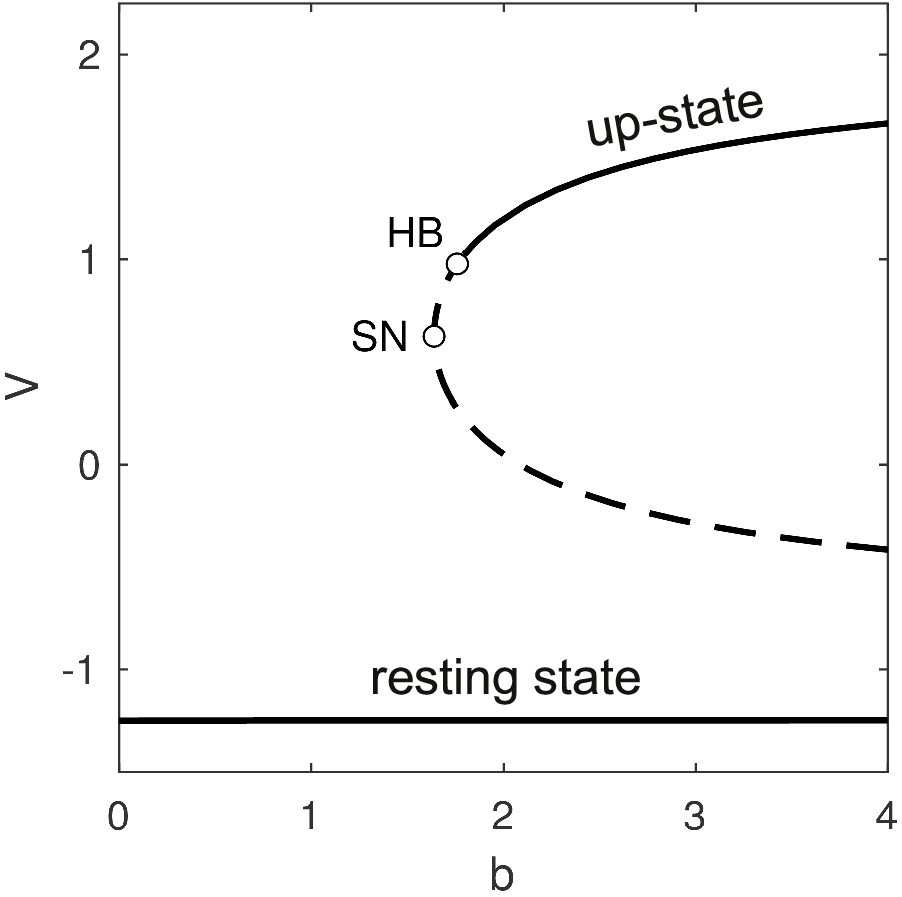
Bifurcations in the steady-states of the single cell. The resting state (*V* = − 1.25) is stable (solid line) for all values of parameter *b* > 0. A pair of unstable fixed points (dashed lines) emerge via a saddle-node (SN) bifurcation at *b*=1.64. The lower branch is the saddle and the upper branch is the node. The node becomes stable via a Hopf bifurcation (HB) at *b*=1.76. The stable up-state co-exists with the stable resting state for *b* ≥ 1.76.

### Mixed medium of normal and non-repolarizing cells

To assess how the presence a limited number of bistable cells might impact the macroscopic electrical properties of heterogeneous cardiac tissue, we constructed a 300 × 100 sheet of cells in which the cells were randomly configured to be either monostable (*b* = 1) or bistable (*b* = 3) in varying proportions. In each case the sheet was stimulated briefly (*I*=2 for 15 ms) at the left-hand boundary to induce a rightward propagating wave (Figure 3A-D). We found that the heterogeneous medium could tolerate a surprisingly large proportion of bistable cells (75%) and still support a functional traveling wave (Figures 3A,B). When that proportion was increased to 85%, the propagating wave left persistent disordered spatiotemporal activity in its wake (Figures 3C,D). In that case, the ectopic activity grew from a small cluster of bistable cells that failed to repolarize in the wake of the traveling wave (Figure 3E). These ‘rogue’ cells remained permanently depolarized and subsequently initiated new propagating waves that ultimately recruited the entire spatial domain. Moreover, those new waves were spatiotemporally disordered like those of the fibrillating state.

**Fig 3.**
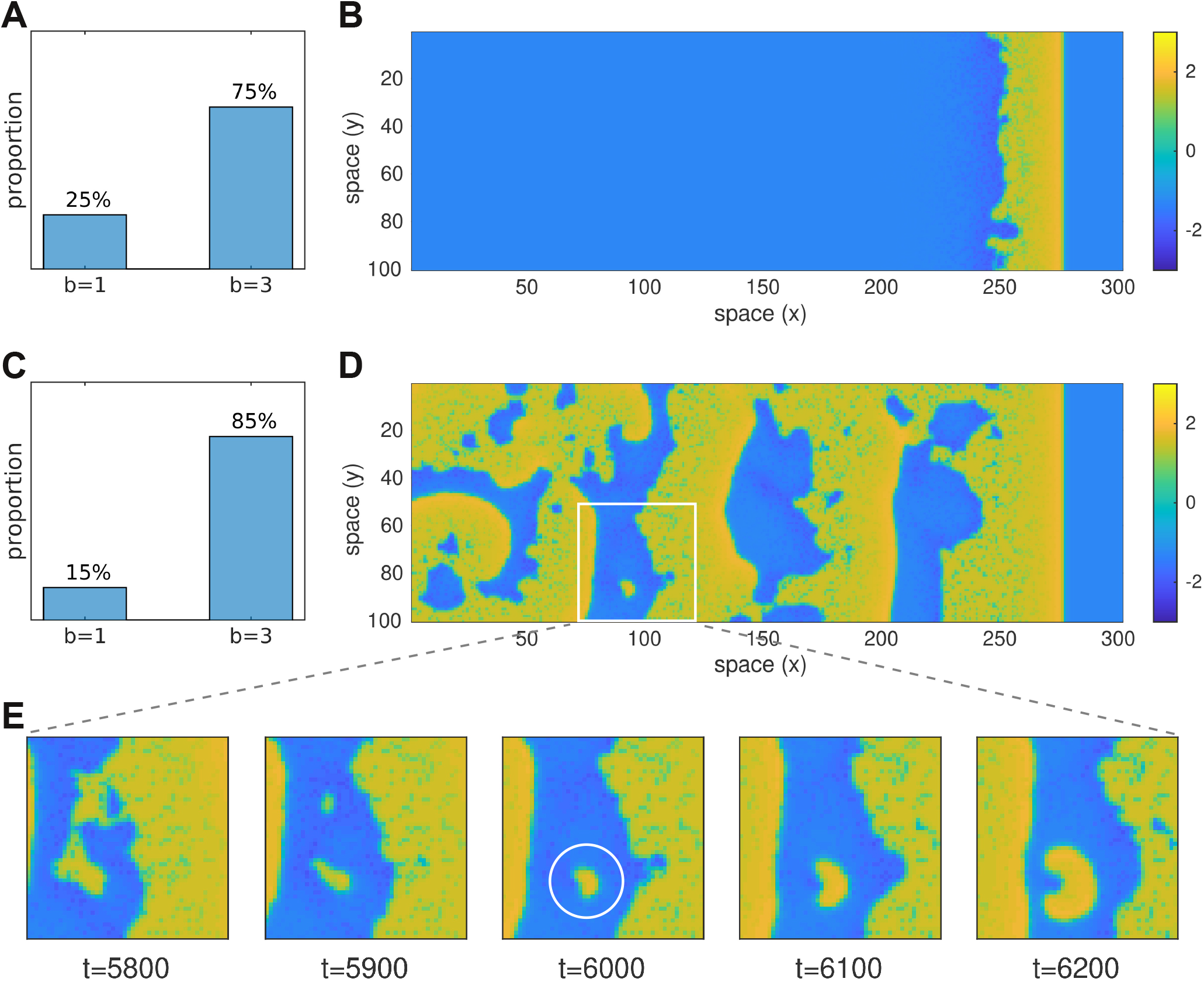
Effect of mixing monostable and bistable cells in the same tissue. Here we compare wave propagation in two simulations of 100 × 300 heterogeneous media where each cell was randomly assigned either *b*=1 (monostable) or *b*=3 (bistable) according to the distributions in panels A and C. A rightward traveling wave was initiated in the resting medium by briefly stimulating the cells at the left hand boundary. Snapshots of the respective simulations (*t*=6000 ms) are shown in panels B and D. The former supports a functional propagating wave whereas the latter leaves self-sustained ectopic activity in its wake. The coupling strength (*c*=0.6) is identical in both cases. Panel E shows successive snapshots of the 50×50 region marked with a white square. It illustrates how the ectopic activity grew from a small cluster of cells that failed to repolarize (white circle). See Supplementary Video S1 for an animated version of this figure.

### Reduced model

To further analyze the conditions under which a bistable cell fails to repolarize when it is embedded in a medium, we constructed a reduced version of our model where a single cell of interest interacts with a hypothetical resting medium (Figure 4A, left). The membrane potential of that cell (*V_o_*) was free to vary while all other cells were fixed at the resting potential (*V_r_* = − 1.25) on the assumption that their membrane potentials were dominated by the medium acting as a global syncitium. Under these assumptions, the gap current flowing into the cell of interest from one neighboring cell is *I = c*^2^(*V_r_ − V_o_*). Since the neighboring cells were all identical, their contributions were lumped together as one equivalent cell (Figure 4A, right). The combined gap current being *I = nc*^2^(*V_r_ − V_o_*) where *n* is the number of neighbors.

**Fig 4.**
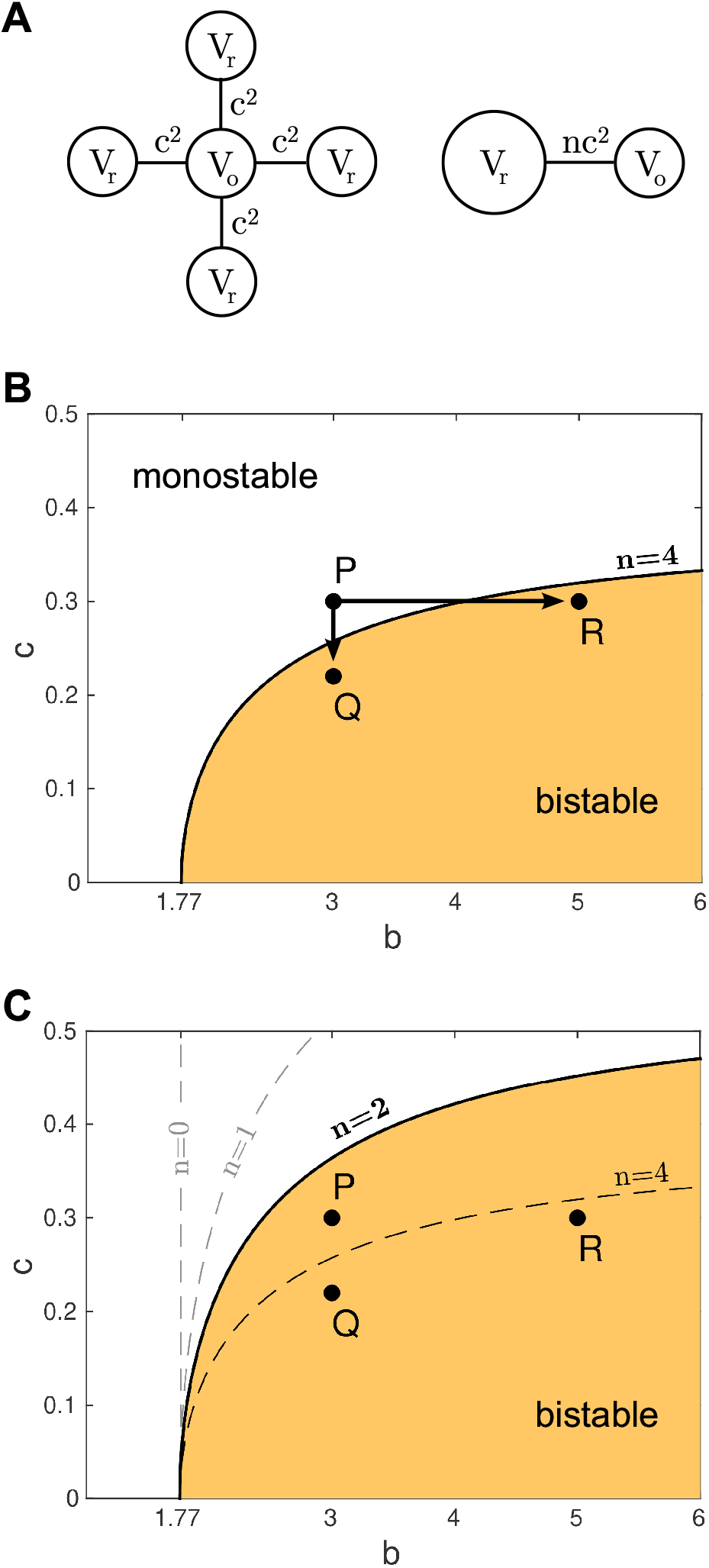
Stability analysis of the reduced model. (A) Schematic of the model. Left: A single bistable cell (*V_o_*) is coupled to n identical cells which are all clamped at rest (*V_r_*). The coupling strength is *c*^2^. Right: The resting cells are lumped together to form the reduced model where the equivalent coupling is *nc*^2^. (B) Stability map for the reduced model with *n*=4 neighbors. The shaded region indicates those configurations of *b* and *c* where the cell operates in the bistable regime. Points *P*={3, 0.3}, *Q*={3, 0.22} and *R*={5, 0.3} are described in the text. (C) Stability map for the case of *n*=2 neighbors. The stability boundaries for *n*=0, *n*=1 and *n*=4 neighbors are included for comparison.

The equations for the reduced model were thus defined as,

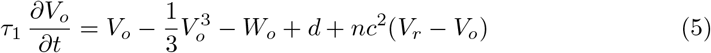

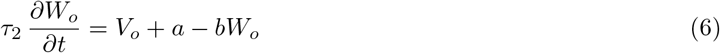

where the parameters are the same as for the single cell equations. Indeed, equations (5–6) have the same basic form as the single cell equations (7–8) hence the bifurcation structure is the same as Figure 2 except that here the location of the Hopf bifurcation (HB) depends on *n* and *c* as well as *b*. We mapped out this critical relationship for the case of *n*=4 by numerically following the Hopf point while allowing *b* and *c* to vary as free parameters. The resulting curve (Figure 4B) describes the stability boundary between the monostable and bistable operating regimes for all configurations of the reduced model. The shaded region indicates those parameter configurations where the cell of interest supports both a stable resting state and a stable up-state, despite the repolarizing influence of the resting medium in which it is embedded. Conversely, the unshaded region indicates those parameter configurations where the cell of interest behaves in a monostable fashion. Cells with those configurations are guaranteed to repolarize when they are embedded in a resting medium even though those with *b* > 1.77 would not do so in isolation.

### Progression of disease

The stability analysis of the reduced model suggests three pathways by which a cell may transition from normal (monostable) behavior — where the cell always repolarizes — to abnormal (bistable) behavior where the cell fails to repolarize. The first pathway involves increasing the intrinsic bistability characteristics of the cell while holding the coupling strength fixed. It is illustrated in Figure 4B by moving the parameter configuration from the monostable regime at *P* to the bistable regime at *R*. In this case, the stability boundary is crossed at *b* ≈ 4. We envisage this pathway as corresponding to some underlying change in the cell’s physiology that impairs its repolarizing currents.

The second pathway involves an overall reduction in the coupling strength between cells while all other aspects of the cell physiology remain unchanged. It is illustrated in Figure 4B by moving from configuration *P* to configuration *Q*. Here the stability boundary is crossed at *c* ≈ 0.26. We envisage this pathway as representing an overall reduction in the conductance of the gap junctions.

The third pathway involves a reduction in the number of neighboring cells. It is illustrated in Figure 4C by the shift in the stability boundary when the number of neighbors is reduced from *n* = 4 to *n* = 2 thus transforming configuration *P* from monostable to bistable. We interpret this particular pathway as highlighting a natural deficit that is faced by cells on tissue boundaries rather than a progressive loss of connectivity over time. It illustrates how cells on tissue boundaries are more susceptible to abnormal behavior than their counterparts in the midfield of the tissue. These boundary effects may explain why the pulmonary vein often appears to be the source of ectopic activity in clinical observations.

### Bistability as a driver of ectopic activity in tissue

Our analysis of the reduced model predicted that a bistable cell in a hypothetical resting media will behave abnormally when the coupling strength is reduced or when it is coupled to fewer neighbors. We tested these predictions in the full model by configuring 10% of the cells in the medium to be intrinsically bistable and the remaining 90% to be intrinsically monostable. We reasoned that this sparse allocation of bistable cells would closely match the reduced model where all neighboring cells were assumed to be at rest. We used a small 40×40 spatial domain so that individual cells could be visualized easily. We also introduced an annulus into the medium to test the prediction that ectopy is more likely to arise at tissue boundaries. The annulus represents a vein or artery in the wall of the heart and was modeled as a region with zero coupling between cells. Cells on the boundary of the annulus had either *n* = 2 or *n* = 3 neighbors depending on the spatial discretization of the local curvature (Figure 5). The spatial domain itself had periodic boundary conditions to eliminate artificial boundary effects.

**Fig 5.**
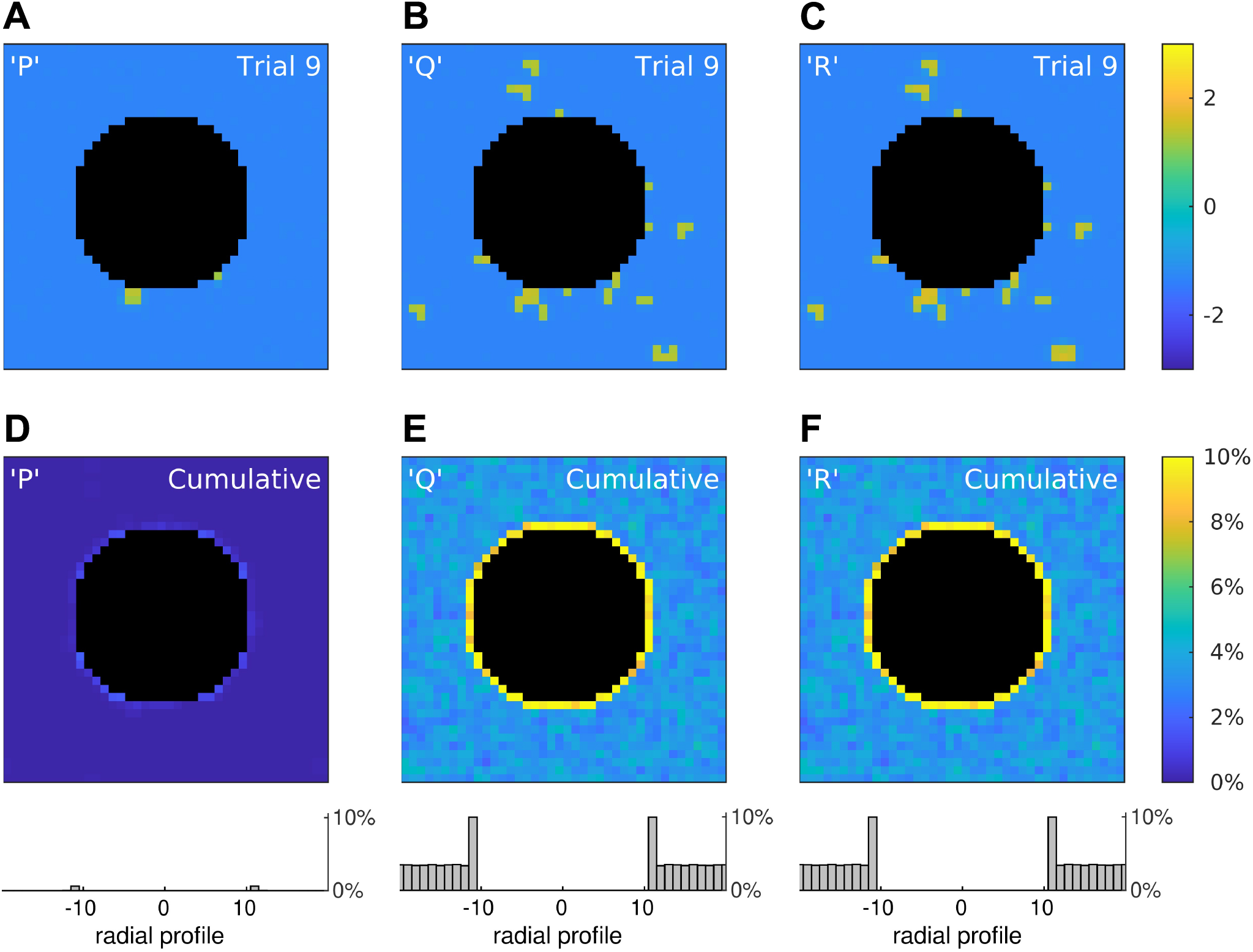
Rogue activity emerges preferentially at tissue boundaries. (A–C) Snapshots of three simulations of a 40×40 medium in which 10% of the cells were configured to points *P, Q* or *R* in the stability map of the reduced model (Figure 4). The snapshots were taken at t = 1000 ms post-onset of a spatially uniform stimulus. The central annulus (black) represents a tissue boundary where cells are absent. The edges of the simulation domain used periodic boundary conditions to eliminate artificial boundary effects. The color scale indicates the membrane potential of the cells. (D–F) Cumulative results for 1000 trials where color is the percentage of cells that remained depolarized in the long term. Histograms (bottom) show the averaged radial profiles. See Supplementary Video S2 for an animated version of this figure.

We simulated the full model under three conditions where the configurations of the bistable cells were taken from points *P, Q, R* on the stability map for the reduced model, respectively. Configurations *Q* and *R* were both predicted to be bistable for *n* = 4 and *n* = 2 neighbors, so those cells were expected to fail to repolarize no matter where they were located in the tissue. Whereas configuration *P* was predicted to be monostable for *n* = 4 neighbors and bistable for *n* = 2 neighbors. Hence those cells were expected to repolarize normally in the midfield of the tissue but not at the boundary of the annulus where the cells have two neighbors. Configuration *P* also straddles the stability boundary for *n* = 3 (not shown) where bistability is marginal. So boundary cells with three neighbors were expected to repolarize normally.

For each configuration, the medium was probed for abnormal repolarization by briefly stimulating it with a spatially-uniform stimulus (*I*=2 for 15 ms) and then observing whether any cells remained depolarized at 1000 ms post-onset of the stimulus. The outcomes of a single trial for each configuration under identical spatial conditions are shown in Figures 5A–C. A survey of 1000 such trials with random spatial conditions is provided in Supplementary Movie S2. Overall, those trials confirm that cells with configurations *Q* and *R* can fail to repolarize at any location, whereas those with configuration *P* are most likely to fail when located at the tissue boundary.

Those findings are quantified by the trial-averages (Figures 5D–F) where color indicates the percentage of trials in which each cell failed to repolarize (*V*>0 at *t*=1000). The histograms (bottom) are the averaged radial profiles of all trials in each condition. They reveal elevated failure rates for cells on the tissue boundary compared to cells in the midfield. In particular, cells on the boundary for configuration *P* failed to repolarize in 0.6% of trials (SE 0.03) while those in the midfield always repolarized. Whereas cells on the boundary for configuration *Q* failed to repolarize in 10.0% of trials (SE 0.12) and those in the midfield failed in 3.4% of trials (SE 0.03). Similarly for configuration *R* where cells on the boundary failed to repolarize in 10.0% of trials (SE 0.12) and those in the midfield failed in 3.5% of trials (SE 0.03). Indeed, the radial profiles in Figures 5E and 5F are virtually identical because configurations *Q* and *R* are equivalent distances from the stability boundary in Figure 4B.

Normalizing the observed failure rates by the density of abnormal cells in the simulated medium (10%) gives adjusted failure rates of 100% for *Q* and *R* cells on the tissue boundary and 35% in the midfield. The adjusted failure rate for *P* cells on the tissue boundary is 6%, although that figure is likely to be an underestimate because many boundary cells had three neighbors instead of two and so were more likely to repolarize. The cells that did fail were predominantly located at the diagonal quadrants of the annulus, as can be seen in Figure 5D. Overall, the simulations in the full model were consistent the predictions of the reduced model.

### Bistability in the context of physiological variability

The analysis above considers a simple binary population of cells that that are either intrinsically monostable (*b*=1) or bistable (*b*=3). While this approach is appropriate for a theoretical analysis of how abnormal cells with bistable characteristics lead to emergent ectopy in coupled tissue, the physiological reality is that cellular properties are not distributed in this manner. Rather, repolarization times of cardiac myocytes are smoothly distributed [15,16], extending to non-repolarizing cells at the extreme long tail of the continuum. To approximate this in our simulations we replaced the sparse binomial distribution used in Figure 5 with a log-normal distribution (*μ, σ*) as shown in Figure 6. The log-normal distribution has a suitably long tail and satisfies *b* > 0 which prevents runaway growth of the recovery variable (Equation 2).

**Fig 6.**
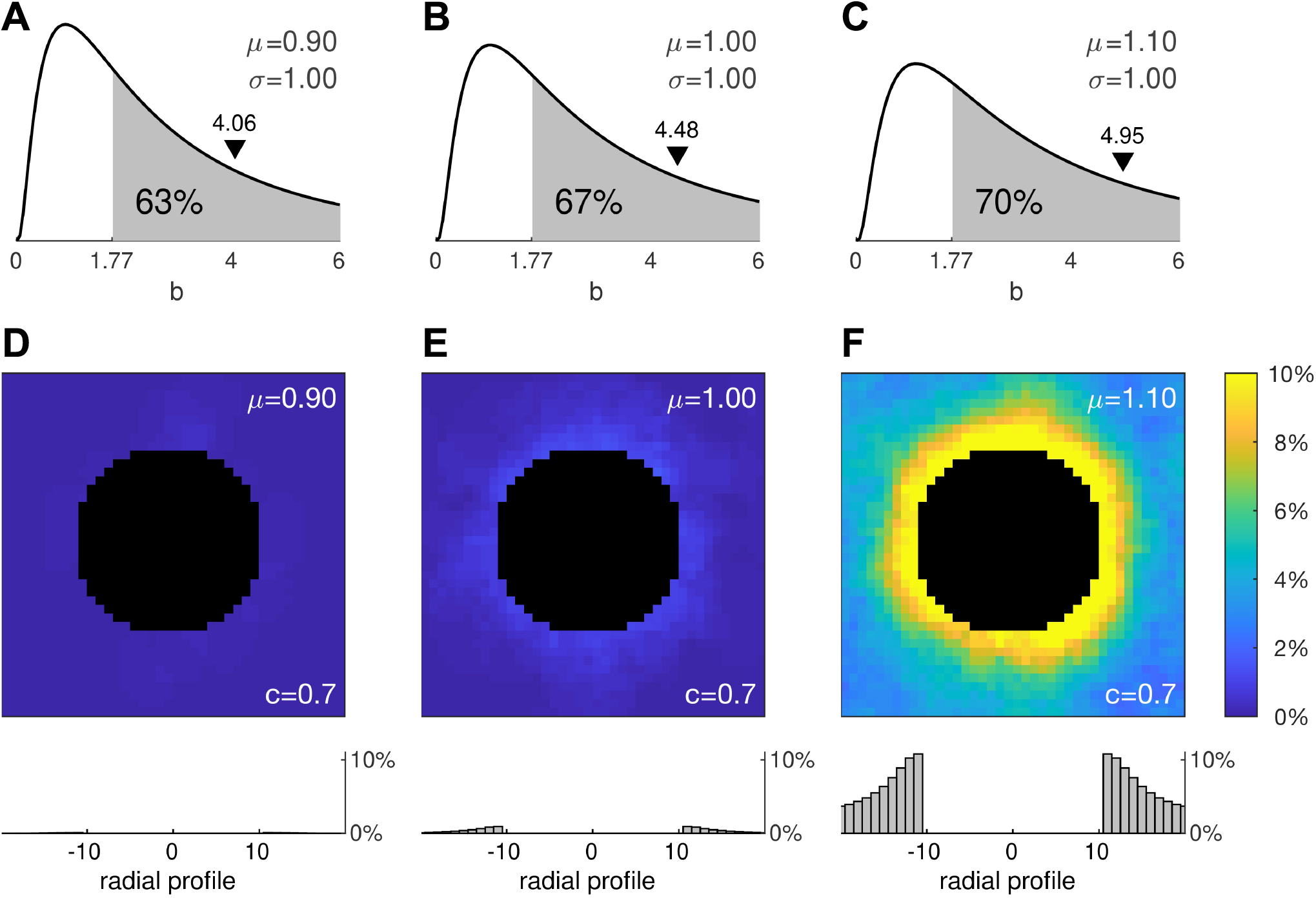
Effect of physiologically realistic distributions of cell heterogeneity. (A–C) Log-normal (*μ, σ*) distributions for *μ*=0.9, *μ*=1.0 and *μ*=1.1 where the second moment is fixed at *σ*=1. The shaded region denotes the area under the curve where *b*>1.77 exceeds the critical value for bistability in the single cell. The black triangle marks the arithmetic mean of the distribution. (D–F) Cumulative results for 1000 stimulation trials using the same protocol as Figure 5. Histograms (bottom) show the averaged radial profiles.

We reasoned that broadening the log-normal distribution would have a similar effect to that of increasing the proportion of bistable cells in the previous simulations. We tested this prediction by manipulating *μ* while holding *σ* = 1 fixed as shown in Figures 6A–C. The shaded regions indicate the proportions of cells in each distribution that are intrinsically bistable (*b*>1.77). Following the same protocol as before, we applied a spatially uniform stimulus to a 40× 40 medium with a central annulus and quantified how many cells remained depolarized at 1000 ms post-stimulus onset. After running a few pilot trials, we fixed the coupling strength at *c* = 0.7.

The trial averages of 1000 simulations in each condition are shown in Figures 6D–F. The first thing that is evident from these simulations is that the tissue can still repolarize normally despite 63% of its cells being bistable. Failure of repolarization does not emerge until that proportion approaches 67% and it is abundantly evident once that proportion reaches 70%. This remarkable tolerance to the presence of abnormal cells is reminiscent of our earlier observations in the densely mixed medium of monostable and bistable cells (Figure 3).

The second thing that is evident is that ectopic activity emerges first at the tissue boundary. Once again, this is consistent with the reduced model, albeit the ectopy emerges at a higher coupling strength than predicted. That is because the densely populated medium allows more cells to reside in the up-state simultaneously and those cells exert no repolarizing influence on one another. Stronger coupling compensates for that loss by amplifying the contributions from the cells that do repolarize.

We next examined how that behavior would manifest in the context of a wave of action potentials propagating across densely heterogeneous tissue containing a natural tissue boundary (Figure 7). In this simulation, the medium was configured with the same parameters (*μ*=1, *σ*=1, *c*=0.7) as per Figure 6E. A rightward traveling wave was initiated in the medium by briefly stimulating the cells on the left-hand side. That wave propagated around the annulus and exited the medium to the right (Figures 7A–E). All cells repolarized normally in the wake of the wave except for a few located on the boundary of the annulus, marked by the white circle in Figure 7G. Those cells subsequently emitted trains of ectopic waves into the surrounding tissue (Figures 7I-M). The simulation is best viewed in Supplementary Movie S3. It demonstrates how sustained fibrillation can be initiated by a small number of cells that fail to repolarize in the wake of a normal propagating wave. It also illustrates how cells at tissue boundaries are more susceptible.

**Fig 7.**
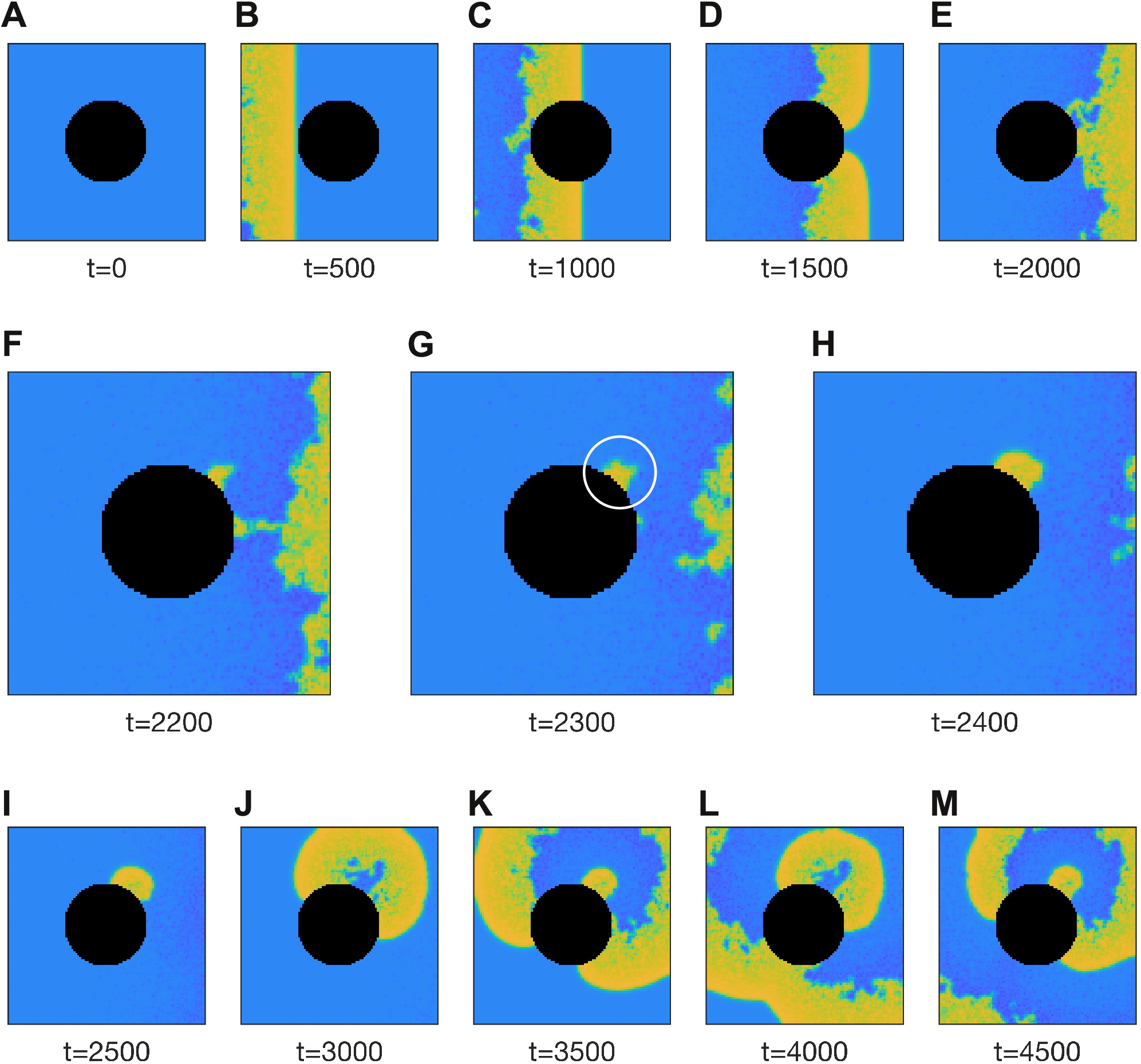
Ectopic activity induced at the tissue boundary by a passing wave. The 100 × 100 medium is briefly stimulated at the left-hand boundary and the ensuing wave propagates to the right. The black region represents the tissue boundary. The white circle in panel G marks the genesis of ectopic activity by cells on the tissue boundary that fail to repolarize. The medium is densely heterogeneous with parameter *b* of each cell drawn randomly from a log-normal distribution (*μ*=1, *σ*=1). The coupling coefficient is *c* = 0.7. See Supplementary Video S3 for an animated version of this figure.

### Progression of Arrhythmic Disease

The observations above provide a framework for understanding how the presence of cells with bistable electrophysiological characteristics may lead to the emergence and progression of arrhythmic disease in the heart. The relationship between the the prevalence of bistability, the degree of inter-cellular electrical coupling, the presence of a tissue boundary and the emergence of disordered tissue level electrophysiology is summarized in Figure 8. In this scheme, the axes represent the pathways for the progression of disease by increasing bistability (left-to-right) and decreasing inter-cellular coupling (top-to-bottom). Each panel is a snapshot of the simulation using the same protocol as Figure 7. The increase in bistability was achieved by increasing the first moment of the log-normal distribution of *b* from *μ* = 0.9 to *μ* = 1.1, exactly as in Figures 6A–C. The inter-cellular coupling was decreased from *c* = 0.8 to *c* = 0.6. The impact of the tissue boundary was encapsulated by the annulus. Overall, the transition from healthy electrical activity (top left) to arrhythmic electrophysiology (bottom right) can be achieved by increasing the prevalence of bistable cells or by decreasing the coupling between cells, or both.

**Fig 8.**
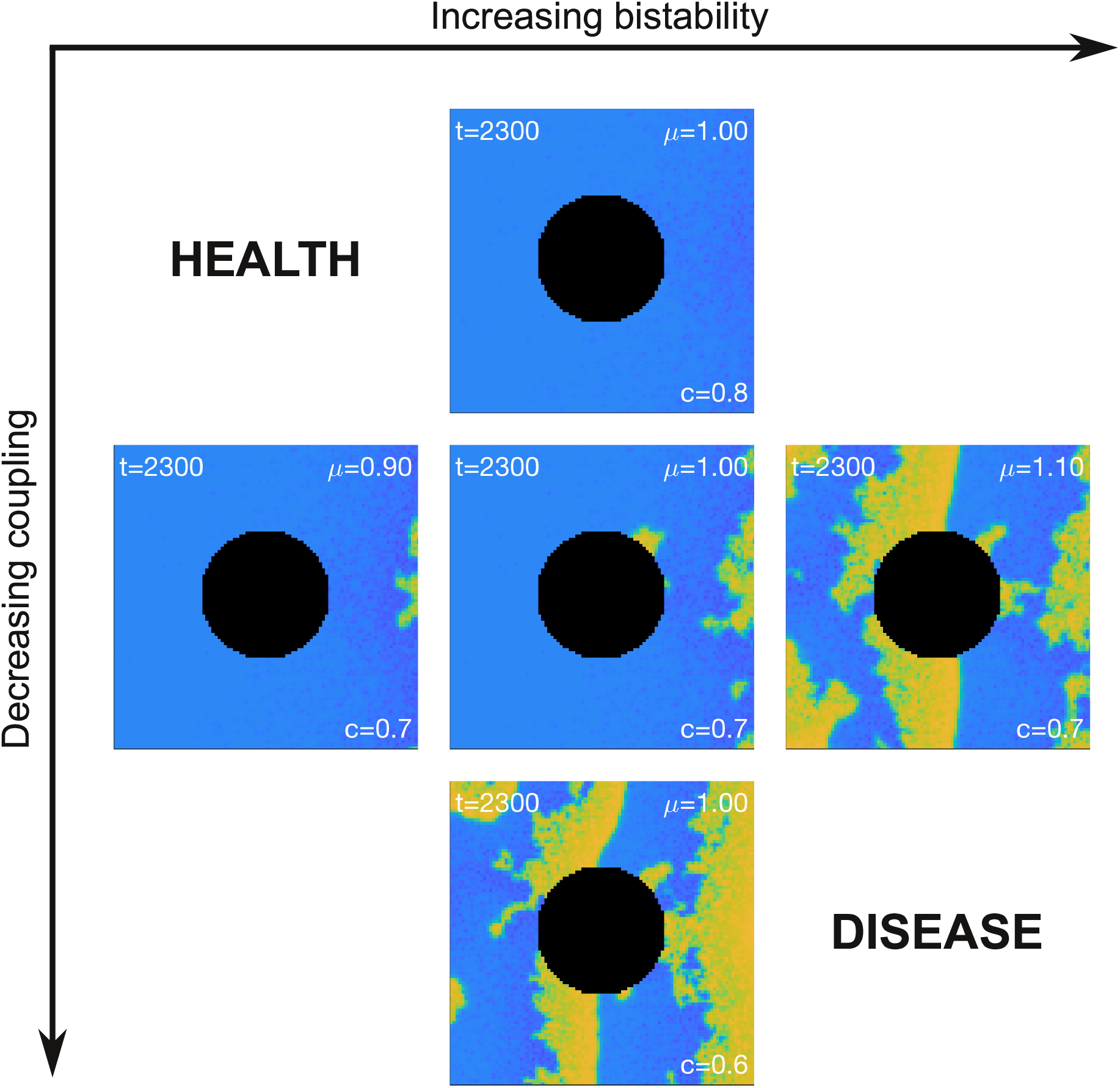
Pathways for the progression of disease. Healthy tissue corresponds to cells with low intrinsic bistability and high coupling (top-left). Disease can progress by increasing bistability or decreasing coupling strength or both. See Supplementary Video S4 for an animated version of this figure.

## Discussion

Recent studies suggest that cardiac myocytes could, under some circumstances, fail to repolarize on their own accord because of inherent bistability in their membrane dynamics [17]. To explore the implications of this phenomenon we modeled cardiac tissue as a simple excitable medium [18,19] containing a mixture of cells with either monostable or bistable membrane dynamics. The medium could tolerate a surprisingly large number of bistable cells and still operate normally because the bistable cells were impelled to repolarize by the normal monostable cells. Nonetheless there was an upper limit to how many bistable cells could be accommodated before normal function was abruptly lost. At that point, the orderly propagation of cardiac waves gave way to disorderly spatiotemporal activity that resembled cardiac fibrillation. Analysis revealed that the propensity to develop fibrillatory activity was determined by a three-way relationship between the expression of bistability within an individual cell, the strength of cell-to-cell coupling in the tissue, and the number of directly coupled neighboring cells. Gradual changes in each of these factors that might occur with progression of cardiac disease will lead to a tipping point in the dynamical behavior consistent with the observation that there is a sudden onset of cardiac arrhythmias in patients with well-established heart disease [2,21].

### Bistability of cardiomyocyte membrane dynamics

In standard physiological preparations, isolated cells that do not repolarize would normally die from calcium toxicity as a result of the calcium channels being held open by the persistent high membrane voltage. Nonetheless, bistable membrane dynamics have been observed in isolated cardiac myocytes under controlled experimental conditions. Kettlewell and colleagues [17] observed them in patch clamped myocytes where the endogenous calcium membrane current was blocked by nifedipine and replaced by a dynamic clamp protocol. Experimental and modeling studies have similarly demonstrated the existence of ultra-long action potentials in cardiac myocytes [13,22,23]. Unlike bistable action potentials, ultra-long action potentials do eventually repolarize but not necessarily before eliciting ectopy. The dynamical theory of EADs by Qu and Chung [22] suggests that ultra-long action potentials correspond to a quasi-equilibrium state at the plateau voltage and that EADs occur when the plateau voltage loses stability through a Hopf bifurcation — evident by a growing oscillation in the membrane potential. The presence of EADs is often regarded as a surrogate marker for pro-arrhythmic behavior.

In our case, the pro-arrhythmic mechanism is not due to the presence of EADs but is instead due to cells that remain in the depolarized state. In the case of a homogeneous bistable medium, the depolarized state recruits the entire domain and never returns to rest (Figure 1F). However the behavior is very different for a heterogeneous medium where the bistable cells are repeatedly driven between the up-state and the resting-state by the action of their neighboring cells. If enough of those neighbors repolarize normally then they can collectively force the bistable cells to operate normally too (Figure 3B). Yet if there are too few normal cells then the regular propagating wave deteriorates into fibrillating activity patterns (Figure 3D) where the individual cells exhibit variable dwell times in either state. As such, they may resemble cells with ultra-long action potentials. Interestingly, the heterogeneous dwell times are reminiscent of early cellular automata studies [12,24] which relied on heterogeneity in refractoriness. In our model, the presence of bistability thus contributes to both the initiation of arrhythmias and the breakdown of regular waves into disorderly spatiotemporal patterns.

### Factors that influence the onset of arrhythmogenesis

In our studies, we considered three mechanisms that modulate the propensity of bistable cells to induce fibrillatory activity in a heterogeneous medium: (i) the degree of bistability in the cell’s membrane potential, (ii) the strength of coupling between cells, and (iii) the number of neighboring cells to which it is electrotonically coupled (Figure 4). Bistability in the generalized Fitzhugh-Nagumo equations (1–2) is governed by the rate at which the recovery variable decays to rest, represented by parameter *b*. Increasing *b* shortens the time window where the recovery variable can actively repolarize the membrane potential. This prolongs the plateau of the action potential until bistability abruptly emerges at the critical value of *b* = 1.77 (Figure 3). At that point, the cell is no longer capable of repolarizing on its own. Nonetheless, it can still be impelled to repolarize by its neighboring cells which drain it of residual current. In doing so, the up-state of the bistable cell is destabilized and the membrane returns to rest under its own dynamics. Even so, this can only happen when the decay rate of the recovery variable is not too large. Otherwise the up-state becomes too strong for the draining currents to overcome. In that case, the cell remains permanently depolarized. The tables are now turned as the ‘rogue’ cell drives current back into the neighboring cells, causing them to depolarize again and again. The cell thus emits spontaneous waves of action potentials into the surrounding tissue which in turn degenerate into fibrillating activity patterns. This path to arrhythmogenesis corresponds to moving from point *P* to point *R* on the stability map (Figure 4B). We suggest that the increasing *b*-value in our modeling studies, would correspond to cells with an increasingly long intrinsic action potential duration [13,17].

The second mechanism of arrhythmogenesis involves a decrease in coupling strength. The role of reduced gap junction coupling in promoting arrhythmogenesis is very well documented in experimental systems [25,26]. A reduction in cell-to-cell coupling diminishes the electrotonic currents that couple cells so relives the dampening effect of neighbouring tissue in limiting the emergence of ectopy (i.e. reduces the so-called sink-source mismatch [9,27]). In our modeling studies, the up-state of the bistable cells gain stability as the neighboring cells lose their influence. Eventually they become permanently depolarized and subsequently emit spontaneous waves into the surrounding tissue. This path to arrhythmogenesis corresponds to moving from point *P* to point *Q* on the stability map (Figure 4B). The arrhythmic behavior would manifest first in those cells with the most extreme expressions of bistability — those with the highest *b*. Subsequent reductions in the coupling strength would see the recruitment of those cells with lower expressions of bistability. The reduction in coupling strength would also exhibit a concomitant reduction in wave propagation speed that could potentially act as a biomarker to empirically distinguish path *P* → *Q* from path *P* → *R* in physiological observations.

The third mechanism promoting arrhythmogenesis involves a reduction in the number of neighbors to which the rogue cell is coupled. In this scenario, the intrinsic bistability of the cell and the conductance of the cell-to-cell coupling are unchanged. The rogue cell merely has fewer neighbors to impel it to repolarize. This path to arrhythmogenesis corresponds to moving the boundary of the stability map (Figure 4C). Such a scenario could arise through progressive tissue death, or through the intrusion of fibrosis into healthy tissue. It also occurs at tissue boundaries where cells naturally have fewer neighbors than their counterparts in the midfield of the tissue. Spontaneous ectopic activity is thus prone to manifest first at tissue boundaries, irrespective of whether it is driven by path *P* → *Q* or *P* → *R*. Such tissue boundary effects have been reported previously (see [28]) and may, for example, contribute to why the cuff of the pulmonary vein is the source of initiation of atrial fibrillation [7].

### Impact of disease progression

The most clinically important cardiac arrhythmias, atrial fibrillation and ventricular fibrillation, are invariably associated with significant underlying heart disease [1,2]. One of the major difficulties with managing these arrhythmias is that despite often decades of progressive heart disease these arrhythmias have an abrupt onset that cannot be predicted in advance [2,21]. The predictions from our modeling of bistable membrane dynamics indicates that this is exactly what one would expect, i.e. the gradual deterioration in cell connectivity, reduction in number of neighbors due to fibrosis, and or increasing extent of bistability can all combine to lead to a sudden tipping point (Figure 8). Our model, however, cannot reproduce the intermittent nature of paroxysmal atrial fibrillation or explain why some arrhythmias spontaneously terminate. This may be due to the lack of action potential duration restitution in the FitzHugh-Nagumo model [29]. Future studies may overcome this limitation by using more realistic cardiac cell models. Such studies would also allow investigation of potential molecular mechanisms underlying fibrillation induced by bistable membrane dynamics. Nonetheless, the causes of fibrillation are multifactorial and different mechanisms are likely to predominate in different patient populations [30]. Bistable action potentials are but one possible cause.

## Methods

### Partial Differential Equations

The partial differential equations (1–2) were transformed into ordinary differential equations by discretizing space using the method of lines [31]. The spatial derivatives were then approximated by the second-order central differences,

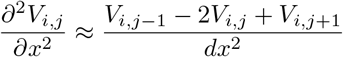

and

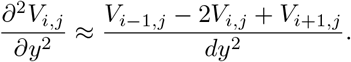

The ordinary differential equations were integrated forward in time using version 2019a of the Brain Dynamics Toolbox [32,33] in conjunction with the Matlab `ode45` solver. The solver error tolerances were `AbsTol=1e-6` and `RelTol=1e-6`.

### Single Cell Equations

The single cell equations,

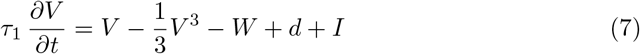

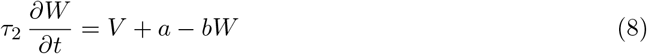

were obtained from equations (1–2) by setting *∂*^2^*V*/*∂x*^2^ = 0. The nullclines (3–4) were obtained by setting *dV/dt* = 0 and *dW/dt* = 0.

### Numerical Continuation

Numerical continuation of the single cell model (Figure 2) was performed using the November 2017 release of the Core Continuation (CoCo) Toolbox [34]. The overall tolerance of the correction algorithm was `TOL=1e-6`. The minimum step size of the continuation algorithm was `h_min=0.01`. The number of discretization intervals for the collocation algorithm was `NTST=20`. Those same tolerances were also used to follow the branch of Hopf points in the stability analysis of the reduced model (Figure 4) with the maximum step size being constrained to `h_max=0.1`.

## Supporting information

**S1 Video. Effect of mixing monostable and bistable cells in the same tissue.** Animated version of Figure 3.

**S2 Video. Rogue activity emerges preferentially at tissue boundaries.** Animated version of Figure 5.

**S3 Video. Ectopic activity induced at the tissue boundary by a passing wave.** Animated version of Figure 7.

**S4 Video. Pathways for the progression of disease.** Animated version of Figure 8.

## Acknowledgments

This work was funded by the Australian government National Health and Medical Research Council grants APP1164518 (AH), APP1182032 (JV) and APP1019693 (JV). We acknowledge support from the Victor Chang Cardiac Research Institute Innovation Centre, funded by the NSW Government.

## Author Contributions

SH, JV and AH contributed to the conception and design of the study. SH wrote the software, performed the simulations and wrote the first draft of the manuscript. JV and AH critically revised the manuscript for intellectual content. All authors contributed to subsequent revisions and they all read and approved the final version.

